# Organic cation transporter 2 contributes to SSRI antidepressant efficacy by controlling tryptophan availability in the brain

**DOI:** 10.1101/2023.02.14.528444

**Authors:** Alejandro Orrico-Sanchez, Bruno P. Guiard, Stella Manta, Jacques Callebert, Jean-Marie Launay, Franck Louis, Antoine Paccard, Carole Gruszczynski, Catalina Betancur, Vincent Vialou, Sophie Gautron

## Abstract

Selective serotonin reuptake inhibitors (SSRI) are common first-line treatments for major depression. However, a significant number of depressed patients do not respond adequately to these pharmacological treatments. In the present preclinical study, we demonstrate that organic cation transporter 2 (OCT2), an atypical monoamine transporter, contributes to the effects of SSRI by regulating the routing of the essential amino acid tryptophan to the brain. Contrarily to wild-type mice, OCT2-invalidated mice failed to respond to prolonged fluoxetine treatment in a chronic depression model induced by corticosterone exposure recapitulating core symptoms of depression, *i*.*e*., anhedonia, social withdrawal, anxiety, and memory impairment. After corticosterone and fluoxetine treatment, the levels of tryptophan and its metabolites serotonin and kynurenine were decreased in the brain of *OCT2* mutant mice compared to wild-type mice and reciprocally tryptophan and kynurenine levels were increased in mutants’ plasma. OCT2 was detected by immunofluorescence in several structures at the blood-cerebrospinal fluid (CSF) or brain-CSF interface. Tryptophan supplementation during fluoxetine treatment increased brain concentrations of tryptophan in wild-type and OCT2 mutant mice, yet more efficiently in WT than in mutants, while discretely increasing 5-HT concentrations. Importantly, tryptophan supplementation improved the sensitivity to fluoxetine treatment of *OCT2* mutant mice, impacting chiefly anhedonia and short-term memory. Western blot analysis showed that glycogen synthase kinase-3β (GSK3β) and mammalian/mechanistic target of rapamycin (mTOR) intracellular signaling was impaired in *OCT2* mutant mice brain after corticosterone and fluoxetine treatment and, conversely, tryptophan supplementation recruited selectively the mTOR protein complex 2. This study provides the first evidence of the physiological relevance of OCT2-mediated tryptophan transport, and its biological consequences on serotonin homeostasis in the brain and SSRI efficacy.

## Introduction

Depression is a widespread and devastating disorder, with up to 17% of the world population at life-time risk of suffering from various forms of this condition [1]. To date, pharmacotherapy can be effective in reducing symptoms and achieving remission, yet a significant proportion of patients suffering from major depressive disorder do not respond adequately to pharmacological treatments. Treatment-resistant depression, defined as failure of at least two consecutive treatments with drugs from distinct pharmacological classes, affects approximately a third of patients [2]. The physiopathological causes underlying this heterogeneity are not fully understood. Current views support the notion that multiple genetic [3-6] and environmental factors such as early life stress [7-9] contribute to shape individual response to treatment. Despite promising leads and current efforts to identify novel treatments [10], selective serotonin reuptake inhibitors (SSRI) and serotonin-norepinephrine reuptake inhibitors remain to date the most prescribed first-line treatment worldwide [11,12]. This class of antidepressants acts principally by blocking high-affinity serotonin (5-HT) and/or norepinephrine (NE) reuptake transporters expressed in aminergic terminals, thereby increasing extracellular concentrations of these monoamines in the brain. Notwithstanding, numerous studies in humans and rodent models argue that the therapeutic effects of SSRI antidepressants also involve other components of the 5-HT circuitry and more broadly neuroadaptive processes occurring over several weeks [13]. The firing of 5-HT neurons in the dorsal raphe nucleus (DRN) directly impacts 5-HT release in projection areas, and the chronic chemogenetic activation of these neurons was shown to induce antidepressant effects in mice [14]. Conversely, during antidepressant treatment, the intrinsic level of somatodendritic 5-HT1A autoreceptors in the DRN and the progressive desensitization of these receptors are believed to tone down the inhibitory action of local 5-HT on 5-HT neuron firing [15,16].

A potential limiting factor in the efficacy of SSRI could be tryptophan routing to the brain. 5-HT in the brain is synthesized from its precursor tryptophan, an essential amino acid supplied by alimentation. 5-HT synthesis is thus highly dependent on tryptophan delivery through specific transporters at the blood-brain barrier [17,18]. Earlier studies in humans and animal models have suggested that prolonged treatments with SSRI require substantial amounts of 5-HT in the brain [19], themselves relying on tryptophan availability. Tryptophan depletion was shown to precipitate relapse in depressed adults in remission treated with SSRI [20-23], and adjunction of the 5-HT precursor 5-hydroxytryptophan has been proposed as a treatment for resistant depression [24]. Mouse models with reduced activity in tryptophan hydroxylase 2 (TPH2), the rate-limiting enzyme in the synthesis of neuronal 5-HT from tryptophan [25], have a deficit in 5-HT levels which is exacerbated by SSRI treatments [19,24,26]. These 5-HT-deficient mice respond poorly to chronic SSRI treatments [26,27] while supplementation with 5-hydroxytryptophan was shown to restore altogether brain 5-HT content [19] and antidepressant response in these animals [26]. It has also been suggested that mutations in the human *TPH2* gene could be associated with poor response to SSRI in small cohorts of patients with depression [28-30].

The mechanisms governing tryptophan penetration in the brain and its influence on antidepressant efficacy are poorly characterized. Organic cation transporters (OCT) are atypical monoamine transporters previously shown to regulate aminergic signaling in the brain and mood-related behaviors such as anxiety, response to stress and antidepressant response [31-35]. Remarkably, one OCT subtype, OCT2, has the capacity to transport tryptophan when expressed in cultured cells [36], in addition to aminergic neurotransmitters. Here, we use a model of chronic depression in mice to test the hypothesis that OCT2 contributes to the long-term effects of the SSRI fluoxetine by regulating tryptophan availability. The findings from this preclinical study reveal a novel role of OCT2 in tryptophan transport to the brain, 5-HT homeostasis and antidepressant efficacy, which adds up with its previously identified function in monoamine clearance within the brain parenchyma [34,37,38].

## MATERIALS AND METHODS

### Animals

*OCT2*^*-/-*^ mice were previously generated by homologous recombination [39]. Heterozygous animals with 10 backcross generations into C57BL6/J were bred to generate wild-type and knockout littermates, which were genotyped as described previously [39]. The behavioral studies were performed during the inactive phase (09:00–13:00) with age-matched (8–16 weeks) male and female mice. Mice were individually housed during corticosterone or corticosterone plus fluoxetine treatment, and collectively housed before basal HLC analysis and immunohistochemistry. To avoid bias, balanced numbers of mice from each group were tested in each daily session. Mice were assigned to groups without randomization. The experimenters were blinded to the treatment of the mice when performing behavioral assessments. Throughout breeding and during the experiments, the mice were fed on a standard diet from Scientific Animal Food and Engineering (A03; Augy, France). Animal care and experiments were conducted in accordance with the European Communities Council Directive for the Care and the Use of Laboratory Animals (2010/63/UE) and approved by the French ethical committee (#5786-2016062207032685).

### Model of chronic depression

To induce a chronic depression-like state, individually housed mice were administered corticosterone in drinking water (35 μg ml-1; Sigma-Aldrich, Darmstadt, Germany) dissolved in 0.45% (wt/vol) hydroxypropyl-ß-cyclodextrin (Sigma-Aldrich) during 10 weeks as previously reported [37,40,41]. Fluoxetine (LKT laboratories, St Paul, MN, USA) was administered intraperitoneally (i.p.) daily at 15 mg/kg/day, a dose effective in previous studies [35,41], during the last 3 weeks of the corticosterone regimens for all experiments except for electrophysiological recording and 5-HT 1A receptor sensitivity experiments, for which a dose of 18 mg/kg was used. Tryptophan supplementation was performed by addition of tryptophan in drinking water (10 mg ml-1, a dose based on previous studies [42]) throughout the fluoxetine treatment. Mice were tested sequentially over 9-day periods for sucrose preference, social interaction, short-term memory in the object location test and anxiety level in the elevated O maze, at basal state, after corticosterone treatment, and after 3 weeks of fluoxetine or fluoxetine plus tryptophan treatment while maintaining corticosterone. Coat state was assessed as a measure of motivation toward self-care. In a first experiment, a group of each genotype (n= 14) was submitted to corticosterone treatment then fluoxetine treatment. Mice of both genotypes (n= 6-9) were removed for western blot analysis after corticosterone and after fluoxetine treatment. In a second experiment, a group of each genotype (n=14-15) was submitted to corticosterone treatment then divided in two groups treated with fluoxetine with or without supplementation. (For details see Supplementary Materials and Methods).

### Tryptophan, 5-HT and kynurenine analysis

Sample collection and biochemical determination of tryptophan, 5-HT and kynurenine concentrations in plasma and tissue extracts were performed by high-performance liquid chromatography (HPLC) with coulometric detection according to procedures previously described. Briefly, brain structures were dissected and homogenized in an ice-cold buffer (0.1 M acetic acid/10 mM sodium metabisulfite/10 mM EDTA/10 mM ascorbic acid) and centrifuged at 22,000 *g* for 20 min at 4°C. Plasma was deproteinized by incubation with two volumes of 0.5 M HClO_4_ supplemented with 0.06 mM ascorbic acid, vortex-mixed and centrifuged at 12,000*g* for 5 min. The collected supernatants were filtered through a 10-kDa membrane (Nanosep, Pall Corp., Port Washington, NY, USA) by centrifugation at 7,000 *g*. Samples were next assessed for monoamine and metabolite content by ultra high-performance liquid chromatography (UPLC Ultimate 3000, Thermo Scientific) and coulometric detection on a Thermo Scientific Hypersil BDS C 18 column including a Hypersil BDS guard column (analytical conditions: isocratic flow at 0.5 mL/min, oven temperature 35°C, Thermo Scientific phase test, analytical cells potentials 100, 250, 450, and 700 mV).

### Immunohistochemistry

Immunohistochemistry was performed as previously described [37]. OCT2 was detected using rabbit polyclonal antibodies (1/500; Agro-Bio, La Ferte St Aubin, France) validated in previous studies [37]. After washing, sections were incubated with Alexa 488-conjugated secondary antibodies (Thermo Scientific, Rockford, IL, USA, Cat# A-11034, RRID:AB 2576217). For double-labeling experiments, antibodies against LAT1 (1/200) from Santa Cruz Biotechnology (CA, USA, Cat# sc-374232, RRID:AB 10988206) were incubated along with OCT2 antibodies and revealed with Alexa 555-conjugated secondary antibodies (Thermo Scientific, Cat# A-31572, RRID:AB_162543).

### Western blots

Whole tissue extracts were prepared from bilateral punches (1–1.5 mm diameter; Miltex, York, PA, USA) of brain regions from adult mice at basal state, treated with corticosterone or with corticosterone plus fluoxetine with or without tryptophan supplementation and submitted to Western blotting as detailed in Supplementary Materials and Methods.

### In vivo electrophysiological recordings of 5-HT neurons in the dorsal raphe

Mice were submitted to recording either at basal state, or after a 3-week treatment with corticosterone, or corticosterone plus fluoxetine, with or with tryptophan supplementation in the drinking water. The day of the recording, they were anaesthetized with chloral hydrate (400 mg/kg i.p.) and placed into a stereotaxic frame. The extracellular recordings of presumed 5-HT neurons in the dorsal raphe (DRN) were performed using single-barreled glass micropipettes (Stoelting Europe, Dublin, Ireland). Their impedance typically ranged between 5 and 8 MΩ. The central barrel used for extracellular unitary recording was filled with 2 M NaCl solution. The glass micropipette was positioned using the following coordinates (in mm from lambda): AP: +1.0 to 1.2; L: 0–0.1; V: 5–7. The 5-HT neurons were then identified using the following criteria: a slow (0.5–2.5 Hz) and regular firing rate and long-duration (2–5 ms) bi- or triphasic extracellular waveform [43]. Details of number of neurons and mice used are given in supplementary table 4.

### 5-HT 1A receptor sensitivity

Mice were submitted to corticosterone treatment as for the model of chronic depression, and fluoxetine was administered for the last 3 weeks in the drinking water at the dose of 18 mg/kg/day. Hypothermia was measured over a period of 30 min after the subcutaneous injection of the 5-HT1A receptor agonist 8-OHDPAT (300 μg/kg; s.c.). Body temperature was assessed using a lubricated rectal probe (BIO-BRET-3, Vitrolles, France) and monitored with a thermometer (Bioseb). Three baseline body temperature measurements at 10 min intervals were taken as a control measure before drug injection, as described by Van der Heyden et al [44], followed by a measurement every 10 min during 1h then two measurements at 90 et 120 min.

### Statistical analysis

PRISM (GraphPad Software, San Diego, CA, USA) was used for statistical calculations. Sample sizes for behavioral and electrophysiology experiments were chosen based on previous reports [35,37,45]. Sucrose preference, social interaction, object location, elevated-O maze, Western blot experiments and HPLC analysis after tryptophan supplementation were analyzed using two-way ANOVA followed by Tukey’s, Sidak’s or Dunnett’s post-hoc tests. Coat state, weight, and 5-HT1A receptor sensitivity were analyzed using repeated two-way ANOVA. HPLC analysis in basal conditions, depression z-scores and western blot experiments after tryptophan supplementation were analyzed using unpaired Student’s *t*-tests. DRN 5-HT neuron firing data did not pass normality and were analyzed using Mann-Whitney tests. Statistical significance was set at P < 0.05.

## RESULTS

### OCT2 is required for the antidepressant effect of fluoxetine in a chronic model of depression

*OCT2-*invalidated mice were previously shown to be insensitive to prolonged treatment with venlafaxine in a chronic depression model [37]. To assess whether this resistance could be extended to another class of antidepressant, we evaluated in the same paradigm the long-term efficacy of fluoxetine, a common antidepressant with a higher selectivity towards the serotonin transporter [46].

To evaluate antidepressant sensitivity, we used a rodent model of chronic depression with strong construct, face and predictive validity [37,40,41] which surpasses acute behavioral despair tests commonly used for antidepressant screening [33,35]. Long-term administration of corticosterone to rodents can induce a panel of behavioral alterations that mimic distinct symptoms of depression. Importantly, these persistent alterations can be improved by long-term, but not acute, antidepressant treatment, as human depression [40,41]. In this model, prolonged corticosterone exposure results in enhanced and comparable levels of circulating corticosterone in *OCT2*□mutants and WT mice [37]. *OCT2* mutant and wild-type (WT) mice were tested in basal conditions, following chronic corticosterone administration and after 3 weeks fluoxetine treatment along with corticosterone, with a sequential battery of tests evaluating anhedonia, social withdrawal, anxiety and cognitive impairment (Fig. 1A). A six week-corticosterone exposure induced robust alterations in sucrose preference, social interaction and anxiety level in the elevated O-maze in both WT and mutant mice (Fig. 1B-E and supplementary Fig. 1-3). Both genotypes showed similar performances in these depression-related behaviors after completion of the corticosterone treatment, as found in a previous study [37] and in another validated depression model, unpredictable chronic mild stress [34]. This contrast with the depressive-like phenotype of *OCT2* mutant mice in resignation tests [37], further illustrating the discrepancies between acute antidepressant-like tests and chronic models of depression [38]. Most behaviors, *i*.*e*., sucrose preference, social interaction and short-term memory, were improved in WT mice but not *OCT2* mutant mice after a three-week treatment with 15 mg/kg fluoxetine (Fig. 1B-D). Similarly, coat state, an indicator of self-grooming behavior and behavioral despair, was altered by corticosterone exposure in both genotypes, but improved by fluoxetine only in WT mice (Fig. 1F). Thus, *OCT2* mutant mice were as sensitive to corticosterone as WT mice but, contrarily to these controls did not show an improvement of their depressive-like phenotype after sustained fluoxetine treatment.

**Fig. 1.**
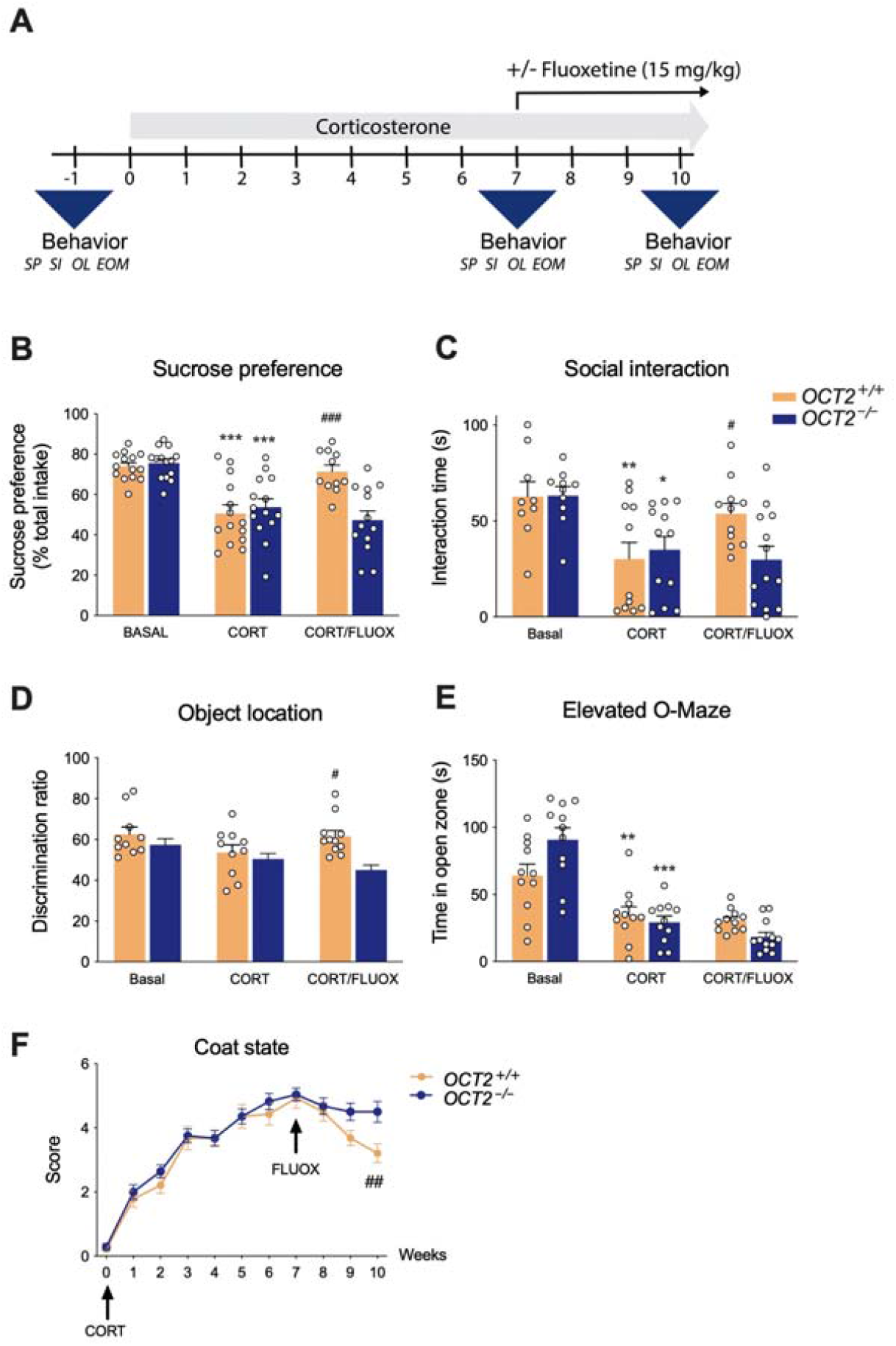
Impaired sensitivity of *OCT2* mutant mice to prolonged fluoxetine treatment in a model of chronic depression. **A** Experimental protocol. Mice were tested sequentially over 9-day periods for sucrose preference (SP), social interaction (SI), short-term memory in the object location test (OLT) and anxiety level in the elevated-O maze (EOM), at basal state, after corticosterone treatment (CORT), and after 3 weeks of fluoxetine plus corticosterone (CORT/FLUOX). Two-way ANOVA analysis (n = 10–14) shows a significant effect of treatment on **(B)** sucrose preference (F2,74= 22.43; P < 0.0001), **(C)** social interaction (F2,60 = 9.3877; P = 0.0003), **(D)** learning and memory in the object location test (F2,56 = 3.863; P = 0.0268) and **(E)** anxiety-like behavior in the elevated O-maze (F2,61 = 43.30; P < 0.0001), and **(F)** of fluoxetine treatment on improving coat state (F1.707,39.82 = 197.7; P < 0.0001). Tukey’s post-hoc tests indicate a significant effect of the corticosterone treatment on sucrose preference, social interaction and time in the open zone of the elevated O-maze in WT (*OCT2*^*+/+*^) and *OCT2* mutant (*OCT2*^*-/-*^) mice (*P < 0.05, **P < 0.01, ***P < 0.001). Three-week fluoxetine treatment significantly increased sucrose preference, social interaction and discrimination ratio in the object location test (Tukey’s post-hoc tests, #P < 0.05, ##P < 0.01, ###P < 0.0001), and improved coat state (Dunnett’s post-hoc tests, ##P < 0.001 week 10 compared to week 7) in WT mice only. Results are given as mean ± s.e.m.

### OCT2 regulates tryptophan, 5-HT and kynurenine levels in the brain and blood

Broadly expressed in the brain [37,38,47], OCT2 participates in 5-HT and NE clearance in aminergic projection areas [37], in complement to the classical monoamine reuptake transporters. Further proof-of concept of its role in monoamine clearance was brought using a selective OCT2 inhibitor designed to penetrate the brain, showing high antidepressant efficacy compared to fluoxetine in a model of chronic depression [35]. On these grounds, the observation that *OCT2* mutant mice were resistant to long-term treatments with the standard antidepressants venlafaxine [37] or fluoxetine (herein) raised a disturbing paradox. Abolition of OCT2-mediated transport resulting from genetic invalidation was expected mainly to increase extracellular 5-HT and NE concentrations in the brain parenchyma, as SSRI, and facilitate antidepressant efficacy. To reconcile these apparently contradictory findings, we speculated that OCT2 could also act at a distinct level from the aminergic synapse, by regulating the availability of the 5-HT precursor tryptophan. To test this possibility, the levels of tryptophan and its other metabolite kynurenine were evaluated in the brain and plasma by high-performance liquid chromatography in *OCT2* mutant mice and their WT littermates. Previous studies evaluating 5-HT content had revealed a marked decrease of tissular levels of this monoamine in *OCT2* mutant mice brain compared to WT [37]. In the present study, *OCT2* deletion was associated with significant reductions in the content of both tryptophan and kynurenine, another tryptophan metabolite, in several brain regions (Fig. 2A), and with increased plasma levels of kynurenine (Fig. 2A), suggesting that OCT2 was involved in the transport of tryptophan to the brain. To gain further evidence supporting this possibility, the eventual presence of OCT2 in regions at the blood-cerebrospinal fluid (CSF) or the brain-CSF interface was explored by immunohistochemistry on mouse brain sections, along with that of the large neutral (or L-type) amino acid transporter LAT1 [18]. Fluorescent immunohistochemistry using previously validated OCT2 antibodies [37] revealed strong OCT2 labeling in the CSF-facing side of ependymal cells bordering the ventricles, also expressing LAT1, shown at the level of the third ventricle in Fig. 2B. OCT2 expression was also found in the arachnoid barrier in the leptomeninges, partially overlapping LAT1, which appeared expressed in addition in the pia mater. Finally, low levels of OCT2 labelling were also found in the choroid plexus and in the subcommissural organ, co-expressed with LAT1 (Fig. 2B). This anatomical data indicates that OCT2 is expressed in several structures regulating metabolites transport into the brain.

**Fig. 2.**
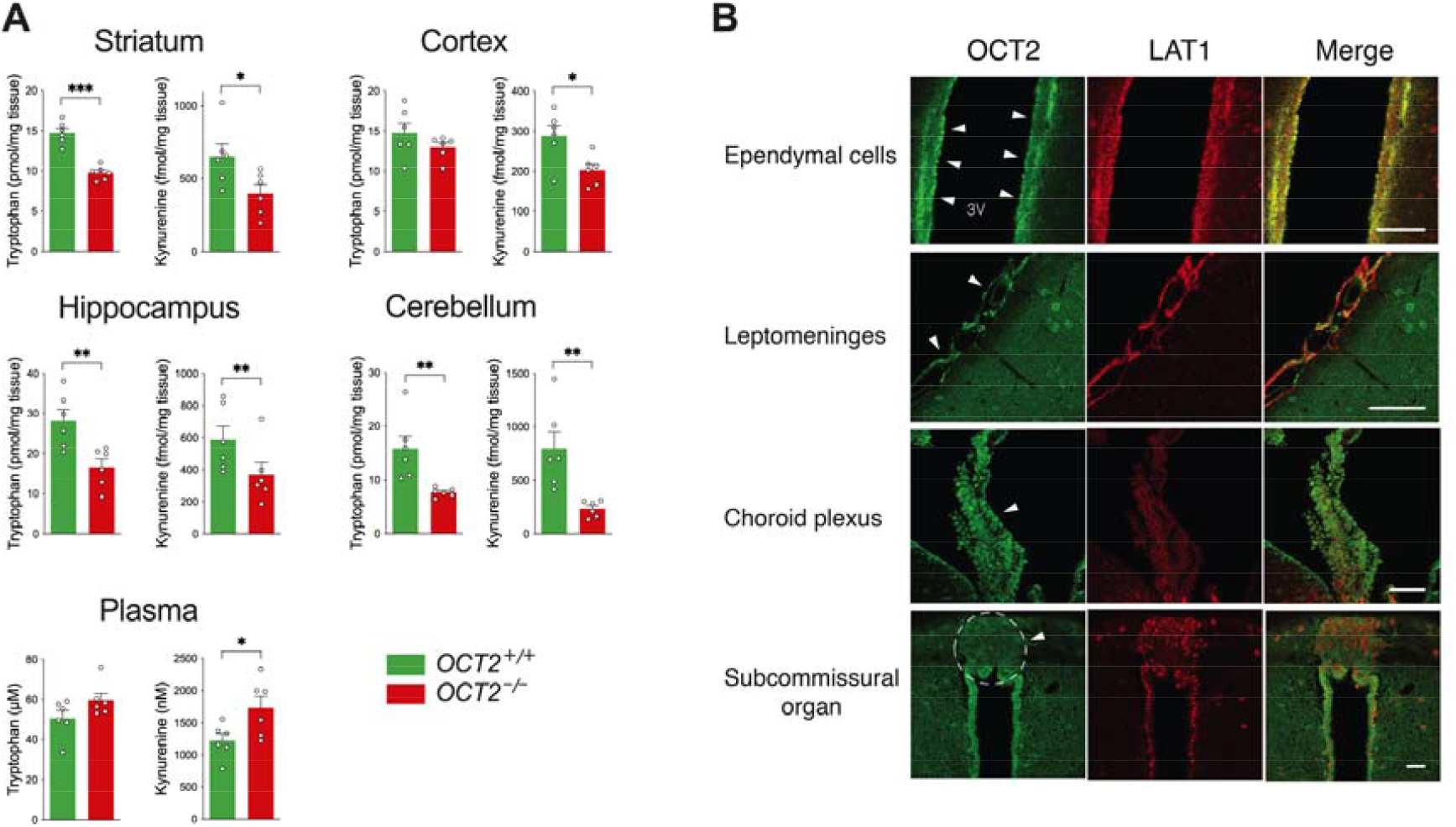
OCT2 activity regulates the levels of tryptophan and its metabolite kynurenine in the brain. **A** High-performance liquid chromatography (HPLC) analysis shows decreased brain levels and increased plasma levels of tryptophan and kynurenine in *OCT2* mutant mice (*OCT2*^*-/-*^) compared to WT mice (*OCT2*^*+/+*^). Two-tailed student’s *t*-tests, unpaired (n = 6; *P <0.05, **P < 0.01). Results are presented as mean ± s.e.m. **B** OCT2 is expressed in structures at the blood-CSF and brain-CSF interface. Immunofluorescent histochemistry of coronal brain sections shows that OCT2 is expressed in the CSF-facing side of ependymal cells boarding the ventricles along with the amino acid transporter LAT1 (shown here at the level of the third ventricle), in the arachnoid barrier in the leptomeninges, partially overlapping LAT1 expression, and at lower levels in the choroid plexus and in the subcommissural organ, also co-expressed with LAT1. Scale bars = 50 μm

### Tryptophan supplementation increases brain tryptophan, 5-HT and kynurenine levels and restores sensitivity to long-term fluoxetine treatment in *OCT2* mutant mice

We next investigated in a series of experiments whether tryptophan availability was responsible for antidepressant resistance in *OCT2* mutant mice. Transport of the 5-HT precursor tryptophan in the brain is ensured mainly by the large neutral amino acid transporters, LAT1 and LAT2. Tryptophan has however a lower affinity for these transporters than their other neutral amino acid substrates [48]. To override this disadvantage of tryptophan over other amino acids for transport across the blood-brain barrier, tryptophan was supplemented in the drinking water of WT and *OCT2* mutant mice during fluoxetine treatment, and the consequences on behavior and tryptophan metabolites content were evaluated.

The effects of tryptophan supplementation on the antidepressant effects of fluoxetine were evaluated in the corticosterone-induced chronic depression model. Sucrose preference (Fig. 3A) and social interaction (Fig. 3B) were robustly altered by corticosterone treatment. Prolonged administration of fluoxetine (15 mg/kg) improved sucrose preference (Fig. 3A), discrimination ratio in the object location test (Fig. 3C), and coat state (Fig. 3D) in WT mice but not in *OCT2* mutant mice, as observed in the first experiment (Fig. 1). Importantly, these three parameters were significantly improved in the *OCT2* mutants when tryptophan supplementation was associated with fluoxetine treatment. Tryptophan supplementation had no effect on body weight in WT or *OCT2* mutant mice (Supplementary Fig. 4). To further evaluate the effects of tryptophan supplementation in *OCT2* mutants, we applied z-normalization across measures of these depression-related variables. Confirming a resistance to the beneficial effects of fluoxetine, the individual z-scores of *OCT2* mutant mice were significantly higher than those of WT mice after administration of the antidepressant (Fig. 3E). Furthermore, the z-scores of *OCT2* mutant mice supplemented with tryptophan during fluoxetine treatment were significantly lower than those of mutant mice treated with fluoxetine alone (Fig. 3E), suggesting an antidepressant effect of this supplementation.

**Fig. 3.**
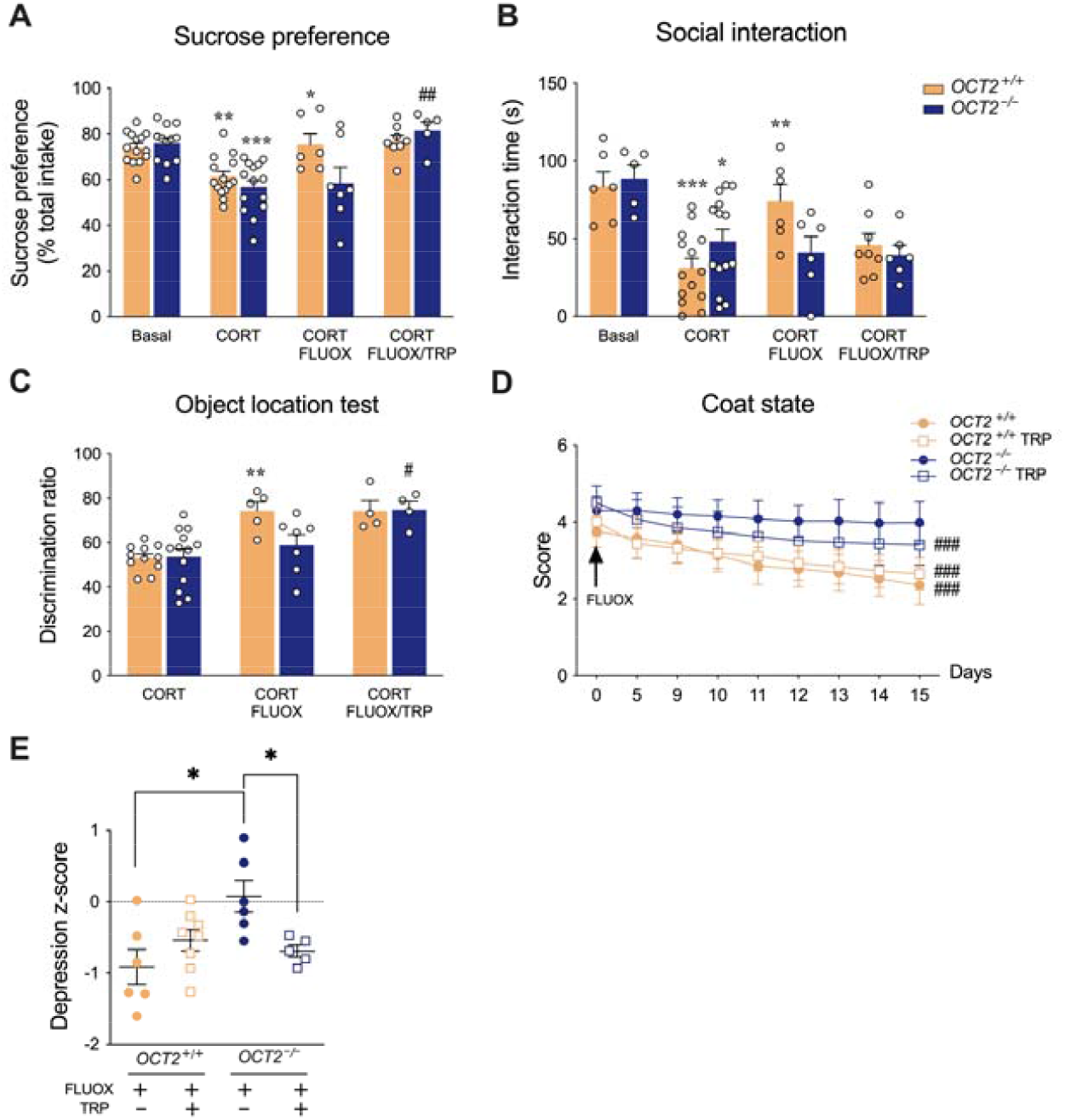
Tryptophan supplementation restores corticosterone-induced depressive-like anomalies of *OCT* mutant mice. Two-way ANOVA analysis (n = 4–14) shows a significant effect of treatment on **(A)** sucrose preference (F3,72 = 16.54; P < 0.0001), **(B)** social interaction (F3,57 = 10.63; P < 0.0001) and **(C)** learning and memory in the object location test (F2,38 = 15.14; P < 0.0001). Tukey’s post-hoc tests indicate significant effects of corticosterone (CORT) treatment (*P < 0.05, **P < 0.01, ***P < 0.001). Three-week fluoxetine treatment (CORT/FLUOX) significantly increased sucrose preference (**A**), social interaction (**B**), and discrimination ratio in the object location test **(C)** in WT mice (*OCT2*^*+/+*^) but not in *OCT2* mutant mice (*OCT2*^*-/-*^), compared with corticosterone alone (*P <0.05, **P < 0.01). Compared to 3-week fluoxetine alone, fluoxetine associated with tryptophan supplementation increased sucrose preference (**A**) and discrimination ratio in the object location test (**C**) in *OCT2* mutant mice. Tukey’s post-hoc test, #P < 0.05, ##P < 0.01. **D** Two-way ANOVA (n=5-8) indicates a main effect of treatment time on improving coat state (F8,176= 56.27; P < 0.0001). Dunnett’s post-hoc tests indicate significant effects of a two-week fluoxetine treatment for WT mice with (CORT/FLUOX/TRP) or without (CORT/FLUOX) tryptophan supplementation and for *OCT2* mutant mice with supplementation (###P < 0.001, day 15 compared to day 0), but not for *OCT2* mutant mice without supplementation. **E** Integrated z-scores across depression-related variables (sucrose preference, social interaction, object location and coat state) after corticosterone followed by fluoxetine treatment (n = 5-8) show a higher depression score in *OCT2* mutants than in WT mice after fluoxetine treatment, and a lower depression score in *OCT2* mutant mice with tryptophan supplementation than without tryptophan supplementation (Two-tailed student’s *t*-tests, unpaired, *P < 0.05). Results are given as mean ± s.e.m.

Next, the impact of tryptophan supplementation on the levels of tryptophan and its metabolites was assessed by HPLC analysis. Tryptophan supplementation during fluoxetine treatment increased significantly the levels of circulating tryptophan and its metabolite kynurenine, both in OCT2 mutant and WT mice (Fig. 4A and supplementary table 3). In addition, tryptophan and kynurenine levels were increased in the plasma of *OCT2* mutant mice compared to WT mice (P < 0.001, supplementary table 3), revealing a role of OCT2 in regulating the blood level of these compounds during fluoxetine treatment. Reciprocally, brain levels of tryptophan, 5-HT and/or kynurenine after fluoxetine treatment were decreased in *OCT2* mutant mice compared to WT mice (Supplementary table 3), confirming the contribution of OCT2 to tryptophan transport into the brain, and its impact on the tissular concentrations of its metabolites. Tryptophan supplementation during fluoxetine treatment induced a significant increase of the concentrations of tryptophan in the three brain areas studied, i.e., striatum, hippocampus, and cortex of WT mice, and in striatum of *OCT2* mutant mice (Fig. 4A and supplementary table 3). This supplementation also increased 5-HT concentrations in the striatum of *OCT2* mutant mice and in the hippocampus of *OCT2* mutant mice (Fig. 4B). 5-HT/tryptophan ratios were increased in OCT2 mutant brain compared to WT treated with fluoxetine with or without supplementation (Supplementary fig. 5), suggesting increased TPH2 activity in the mutants [25]. Thus, in these experimental conditions, tryptophan deficiency in *OCT2* mutants appeared to be moderately and variably compensated by tryptophan supplementation through diet, depending on the brain area considered. In contrast, we detected few significant effects of this supplementation on the brain concentrations of kynurenine, which were more heterogenous (Fig. 4B and supplementary fig. 5).

**Fig. 4.**
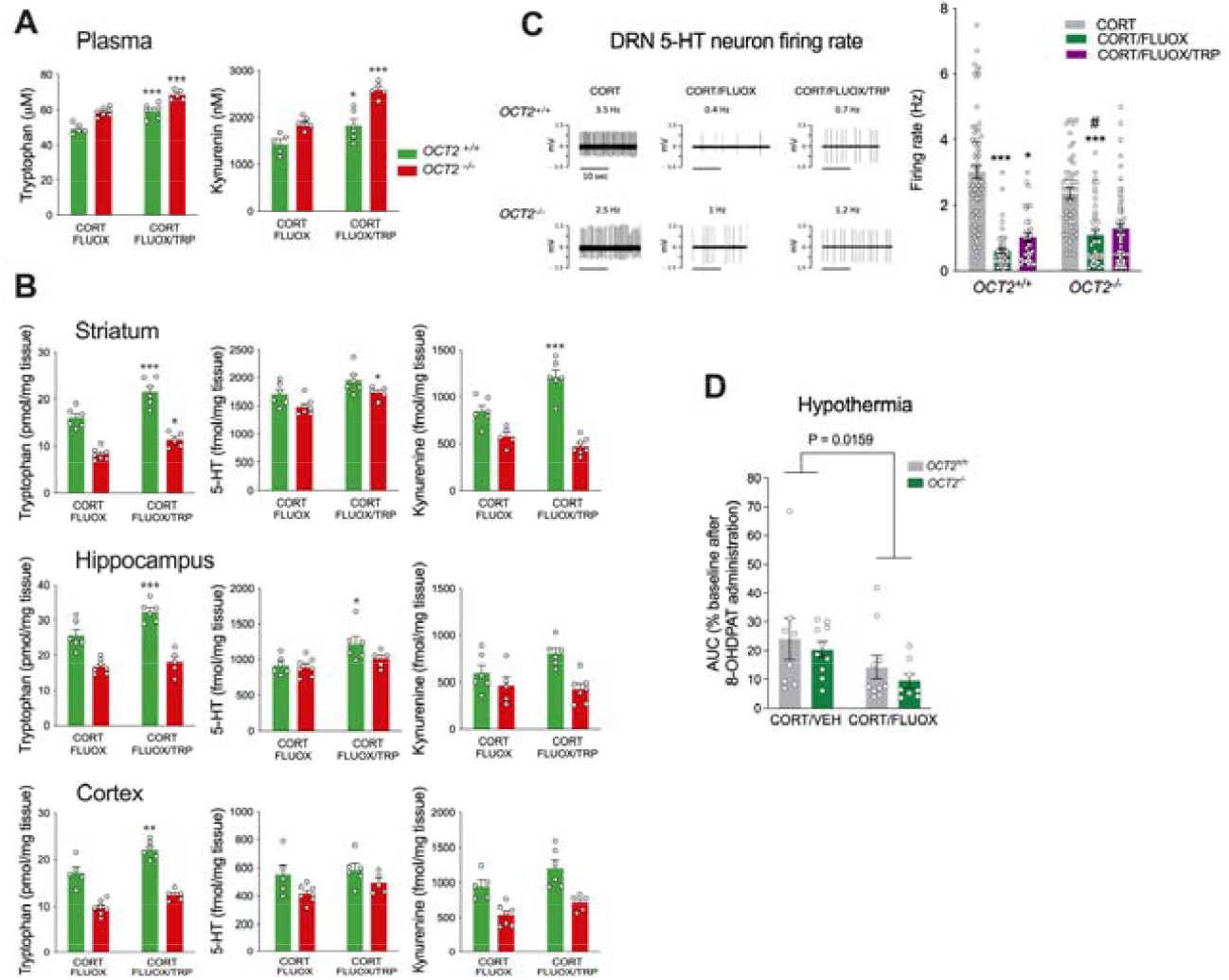
Tryptophan supplementation affects differentially brain tryptophan and kynurenine levels and 5-HT signaling in WT and *OCT2* mutant mice. **A** High-performance liquid chromatography (HPLC) analysis shows increased levels of circulating tryptophan and kynurenine in *OCT2* mutant and WT mice after tryptophan supplementation (CORT/FLUOX/TRP) compared to fluoxetine alone (CORT/FLUOX). Two-way ANOVA analysis (n = 5–7) shows a significant effect of tryptophan supplementation on the levels of plasma tryptophan (F1,19 = 52.22; P < 0.0001) and kynurenine (F1,19 = 38,29; P < 0.0001), and of genotype on the levels of plasma tryptophan (F1,19 = 46.89; P < 0.0001) and kynurenine (F1,19 = 44,64; P < 0.0001). Tukey’s post-hoc tests indicate increased levels of circulating tryptophan and kynurenine in both WT (*OCT2*^*+/+*^) and *OCT2* mutant (*OCT2*^*-/-*^) mice after tryptophan supplementation (*P < 0.05, ***P < 0.001). Main effects detailed in Supplementary table 3). **B** HPLC analysis shows increased levels of tryptophan, 5-HT and kynurenine in several brain regions of WT and *OCT2* mutant mice after tryptophan supplementation. Two-way ANOVA analysis (n = 5–7) show a significant effect of tryptophan supplementation and genotype on the levels of tryptophan in striatum, hippocampus and cortex; of tryptophan supplementation on the level of 5-HT in striatum and hippocampus and on the level of kynurenine in striatum and cortex; and of genotype on the level of 5-HT in striatum, hippocampus and cortex. Main effects detailed in Supplementary table 3. Sidak’s post-hoc tests indicate increased levels after tryptophan supplementation of tryptophan in striatum, hippocampus and cortex of WT mice, and in striatum of *OCT2* mutant mice; of 5-HT in striatum of *OCT2* mutant mice and hippocampus of WT mice; of kynurenine in striatum of WT mice (*P < 0.05, **P < 0.01, ***P < 0.001). Results are presented as mean ± s.e.m.). **C** In vivo electrophysiological recordings of 5-HT neurons in the dorsal raphe nucleus (DRN). Left, typical recordings of DRN 5-HT neurons in the different experimental conditions. Action potentials show the well-characterized regular discharge of 5-HT neurons (scale bar=10 seconds); right, frequency (Hz) of 5-HT neurons firing recorded in the DRN of WT (*OCT2*^*+/+*^) and *OCT2* mutant (*OCT2*^*-/-*^) mice exposed to corticosterone (CORT) alone or in presence of fluoxetine (CORT/FLUOX) or to the combination of fluoxetine plus tryptophan supplementation (CORT/FLUOX/TRP); mean ± s.e.m. Mann-Whitney tests (n = 34–65, 4-7 mice) indicate a significant decrease in firing activity after fluoxetine treatment in corticosterone-treated WT and *OCT2* mutant mice (***P<0.001). Firing after fluoxetine treatment was significantly lower in WT compared to mutant mice (#P < 0.05). Tryptophan supplementation increased 5-HT neurons firing in WT (*P < 0.05) but not *OCT2* mutant mice. **D** 5-HT1A receptor sensitivity assessed using the 8-OHDPAT-induced hypothermia test. Data are means ± s.e.m. of the area under the curve (AUC) representing overall decrease in body temperature (°C) in WT (*OCT2*^*+/+*^) and *OCT2* mutant (*OCT2*^*-/-*^) mice exposed to corticosterone alone (CORT/VEH) or in the presence of fluoxetine (CORT/FLUOX). Two-way ANOVA (n = 8–10) shows an effect of genotype (F1,33 = 6.460; P = 0.0159) but no significant effects of fluoxetine treatment (F1,33 = 2.019; P = 0.16547) or treatment*genotype interaction (F1,33 = 0.3094; P = 0.5818) but on 8-OHDPAT sensitivity. Sidak’s post-hoc tests show no differences between groups.

To further characterize the impact of OCT2 invalidation and tryptophan supplementation, we examined the activity of dorsal raphe nuclei (DRN) 5-HT neurons in *OCT2* mutant mice and their WT littermates after administration of corticosterone alone or in combination with fluoxetine, with or without tryptophan supplementation. Mann-Whitney analysis indicated a comparable firing rate of DR 5-HT neuron in OCT2 mutant mice (1.63 ± 0.13; n= 41; 4 mice) than in WT controls (2.08 ± 0.02; n= 34; 4 mice) at basal state (p= 0.057). A three-week fluoxetine treatment significantly lowered the firing rate of DRN 5-HT neurons in corticosterone-treated WT and *OCT2* mutant mice, compared to corticosterone alone (Fig. 4C). This result is compatible with a 5-HT1A receptor-mediated autoinhibition of DRN 5-HT neuronal activity, a well-documented consequence of increased extracellular 5-HT concentration at the proximity of the 5-HT cell bodies occurring during SSRI administration [16]. This analysis also revealed a higher firing activity in *OCT2* mutant mice compared to WT after fluoxetine treatment, while tryptophan supplementation significantly increased firing in WT but not *OCT2* mutant mice. Measure of 8-OHDPAT-induced hypothermia showed a main effect of fluoxetine treatment, but no significant differences in 5-HT1A autoreceptors sensitivity between genotypes treated with corticosterone alone or in association with fluoxetine (Fig. 4D). Taken together, these different observations indicate that OCT2 modulates 5-HT homeostasis and DRN 5-HT neuronal activity in the brain during fluoxetine administration.

### GSK3β and mTOR intracellular signaling is impaired in *OCT2* mutant brain after fluoxetine treatment, and tryptophan supplementation selectively recruits the mTOR complex 2

Intracellular signaling pathways in the brain involving ERK1/2, GSK3β and mammalian/mechanistic target of rapamycin (mTOR) have been implicated in depression in humans [49-51] and in antidepressant response in animal models [52-54]. We found previously that the activity of these three pathways (ERK1/2, GSK3β and mTOR) is profoundly altered in the hippocampus and prefrontal cortex by a prolonged corticosterone treatment in mice, and that these anomalies can be corrected by a long-term treatment with fluoxetine [35]. The prefrontal cortex and limbic regions including the hippocampus are highly involved in the cognitive and emotional aspects of mood disorders [55,56], acting as hubs integrating the various neurochemical, synaptic and trophic signals that lead to a depressive state. Thus, evaluating intracellular signaling in these brain areas may allow monitoring the general impact of pro- or anti-depressive conditions in animal models.

To gain further insight on the impact of OCT2 activity and tryptophan supplementation on ERK1/2, GSK3β and mTOR intracellular signaling in the brain, we investigated the activation state of these pathways in the dorsal hippocampus and prefrontal cortex of WT and *OCT2* mutant mice by quantitative Western blot analysis. As found previously in WT mice [35], the phosphorylation state of ERK1/2 and Akt at Ser-473, evaluated by the phosphorylated over non-phosphorylated ratios of these proteins, and of the levels of phosphorylated p70S6 were significantly decreased after corticosterone treatment in the hippocampus and the prefrontal cortex of WT and *OCT2* mutant mice, indicating an inhibition of the ERK1/2, TOR complex 2 (TORC2) and TORC complex 1 (TORC1) signaling, respectively (Fig. 5A). Additionally, as expected from our earlier study [35], phosphorylation of GSK3β at Ser-9 (Fig. 5A) was decreased, indicating activation of this pathway. In WT mice brain, most of these alterations in activity induced by chronic corticosterone were reversed after a three-week treatment with fluoxetine (Fig. 5A), sometimes raising to activation states exceeding that of basal state. In contrast, fluoxetine treatment reversed only partially these alterations in *OCT2* mutant mice brain. Notably, ERK1/2 activity was increased in the hippocampus of both genotypes by fluoxetine treatment, but the phosphorylation state of Akt at Ser-473 and of GSK3β at Ser-9 were upregulated in this brain area in WT but not *OCT2* mutant mice (Fig. 5A). This indicated that the regulation of GSK3β and TORC2 signaling occurring during fluoxetine administration was impaired in the *OCT2* mutants’ hippocampus. In the prefrontal cortex of WT and OCT2 mutant mice, fluoxetine treatment increased the phosphorylation state of ERK1/2 and GSK3β at Ser-9. Of note, although not statistically significant, the phosphorylation state of Akt at Ser-473 in WT cortex showed a tendency to increase after fluoxetine treatment, contrarily to *OCT2* mutants, pointing to potential anomalies of TORC2 signaling also in this brain area in the mutants. Our results show that mTOR and GSK3β signaling was profoundly altered in the chronic depression model used in this study, and that fluoxetine reversed in part these alterations in normal mice but not in *OCT2* mutants. In a final step, we questioned whether modifications of these signaling pathways were associated with the improvement by tryptophan supplementation of depressive-like behavior in *OCT2* mutants. Tryptophan supplementation increased phosphorylation of Akt at Ser-473, but not p70S6 nor GSK3β at Ser-9 in the dorsal hippocampus and the prefrontal cortex of *OCT2* mutant mice (Fig. 5B), revealing the recruitment of the mTOR protein complex 2 during supplementation.

**Fig. 5.**
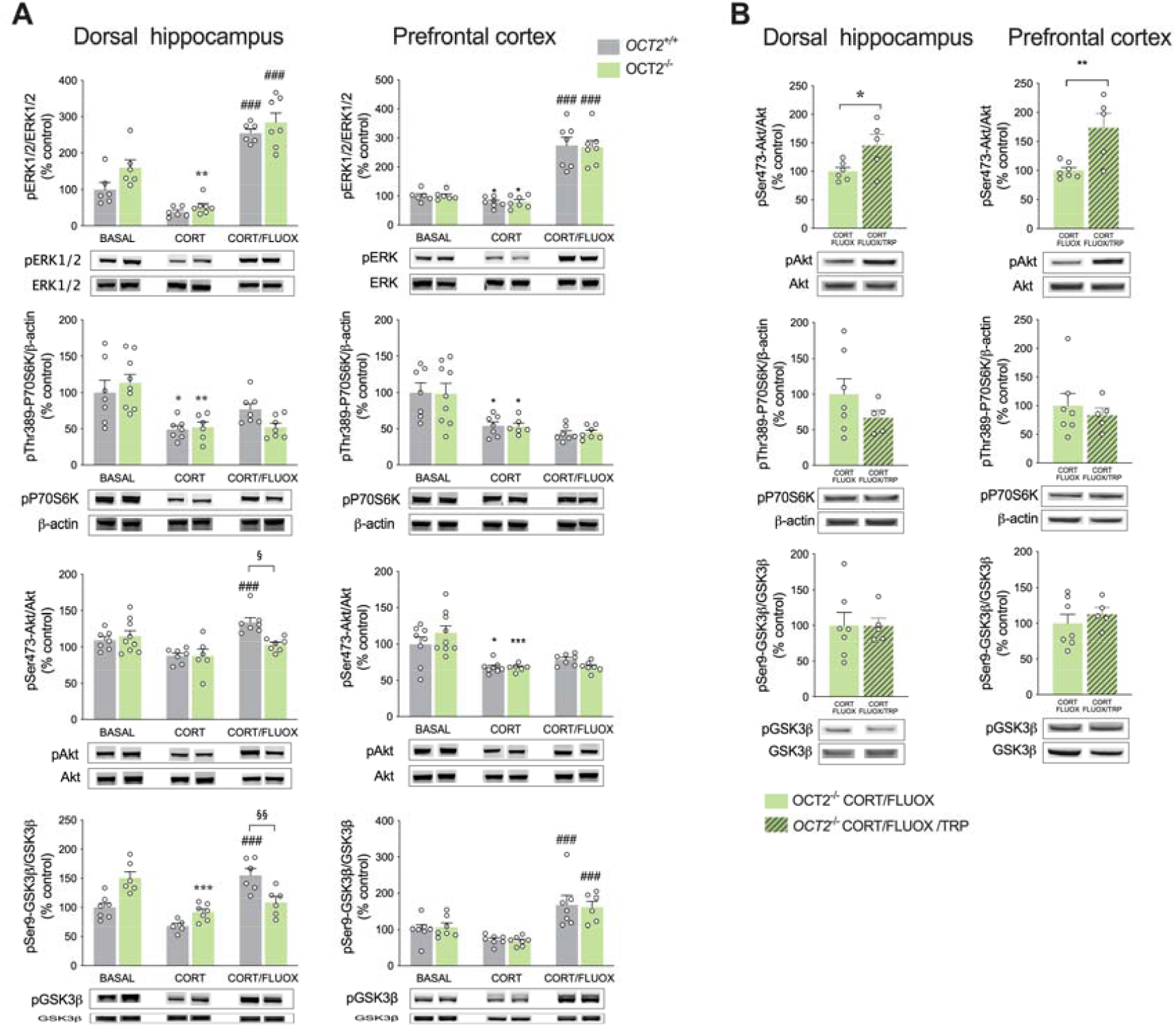
GSK3β and mTOR intracellular signaling is impaired in *OCT2* mutant brain after fluoxetine treatment, and tryptophan supplementation increases selectively mTORC2 signaling in mutant brain. **A** Quantitative Western blot analysis showed alterations of extracellular-signal regulated kinase1/2 (ERK1/2), glycogen synthase kinase-3β (GSK3β) and mammalian target of rapamycin (mTOR) signaling after chronic corticosterone administration (CORT) and fluoxetine treatment (FLUOX). Two-way ANOVA (n = 6–9) show main effects of treatment on the phosphorylation state of ERK1/2 (F2,32 = 87.20; P < 0.0001), on the levels of phosphorylated p70S6 (F2,37 = 16.05; P < 0.0001), and on phosphorylation state of Ser473 Akt (F2,37 = 12.06; P < 0.0001) and Ser9 GSK3β (F2,31 = 21.14; P < 0.0001) in the dorsal hippocampus of WT (*OCT2*^*+/+*^) and *OCT2* (*OCT2*^*-/-*^) mice; on the phosphorylation state of ERK1/2 (F2,34 = 78.72; P < 0.0001), on the levels of phosphorylated p70S6 (F2,37 = 20.12; P < 0.001), and on the phosphorylation state of Ser473 Akt (F2, 39 = 21.75; P < 0.0001) and Ser9 GSK3β (F2,35 = 22.43; P < 0.0001) in the prefrontal cortex (PFC) of WT and OCT2 mice. Tukey’s post-hoc tests show significant differences between untreated (BASAL) and corticosterone-treated (CORT) groups (*P < 0.05, **P < 0.01, ***P < 0.001). Fluoxetine treatment (CORT/FLUOX) increased the phosphorylation state of ERK1/2 in the dorsal hippocampus of WT and *OCT2* mutant mice, and the phosphorylation state of Ser473 Akt and Ser9 GSK3β in the dorsal hippocampus of WT but not *OCT2* mutant mice. Fluoxetine treatment increased the phosphorylation state of ERK1/2 and Ser9 GSK3β in the PFC of both WT and *OCT2* mutant mice. Tukey’s post-hoc tests show significant differences after fluoxetine treatment compared with corticosterone-treated groups (###P < 0.001) and between WT and mutant mice after fluoxetine treatment (§P < 0.05, §§P < 0.01). Results are given as mean of phosphorylated over non phosphorylated protein ratio ± s.e.m, or for p70S6 as mean of phosphorylated protein over β-actin ratio ± s.e.m. **B** Phosphorylation state of Ser473 Akt reflecting TORC2 activation was significantly increased in the PFC and dorsal hippocampus of *OCT2* mice when tryptophan (CORT/FLUOX/TRP) was administered during fluoxetine treatment compared with fluoxetine alone (CORT/FLUOX). Two-tailed student’s *t*-tests, unpaired (n = 5–6; *P < 0.05, **P < 0.01). Results are presented as mean ± s.e.m.

## DISCUSSION

Understanding the heterogeneous etiopathology of major depressive disorder may allow to delineate subsets of depressed patients, which could influence treatment outcome [57]. The present preclinical study reveals a main role of OCT2 in SSRI efficacy through its role in supplying tryptophan to the brain. Prolonged treatments with SSRI require high levels of 5-HT in the brain, which may be restricted by the availability of its precursor tryptophan [58]. The essential amino acid tryptophan is imported into the brain by specific transporters at the blood-brain barrier, notably by the L-type amino acid transporters (LAT) [17,18]. Due to their variable affinity, large neutral amino acids compete for transport at the LAT, with aromatic amino acids such as tryptophan being at a disadvantage compared to other amino acids substrates [48,59]. Previous functional in vitro studies also suggested that tryptophan could be an endogenous substrate of OCT2 [36], but the importance of this transport activity for brain function had not yet been documented. We propose that in conditions of relatively low amino acid supply such as the standard mouse diet, this unfavorable competition of tryptophan with other large amino acids for LAT increases the need for alternate tryptophan transport systems such as OCT2 to ensure sufficient tryptophan levels in the brain. Our results show that OCT2 is expressed in certain structures which may regulate metabolite transport to and from the brain, with a distribution complementing or overlapping LAT1. Fluorescent immunohistochemistry showed the presence of OCT2 at the CSF-facing side of the ependymal cells bordering the ventricles, which form a partial barrier regulating the transport of molecules at the ventricle-parenchyma interface [60,61]. This transporter was also found in the arachnoid membrane (the blood-arachnoid barrier) which isolates CSF from circulating blood [62,63]; at a lower level in the choroid plexus, which controls the flow of metabolites at the blood-CSF interface [64]; and in subcommissural organ secretory cells, which establish a direct contact with CSF and local capillaries [65]. The exact contribution of these structures to tryptophan transport is yet to be determined. In addition, although fluoxetine poorly interacts with OCT2 [66,67], it cannot be excluded that OCT2 deficiency could influence indirectly the clearance, routing to the brain or metabolism of this antidepressant. In spite of these limitations, OCT2-mediated transport of tryptophan across barrier structures is strongly supported by our HPLC analysis. Compared to WT littermates, naive *OCT2* mutant mice showed significant reductions in brain levels of 5-HT [37], tryptophan and kynurenine (herein). These differences were also found after fluoxetine treatment in the corticosterone-induced chronic depression model, with concentrations of these compounds significantly decreased in mutant mice brain compared to WT mice and, conversely, increased in the plasma of mutants, emphasizing the contribution of OCT2-mediated transport in modulating brain levels of these metabolites. Additionally, TRP supplementation increased brain content of tryptophan to a greater extent in WT than in mutant mice, further underscoring the importance of OCT2 for tryptophan transport. However, the differences between genotypes in 5-HT levels were more discreet than those of tryptophan. This could be the consequence of increased TPH2 activity in *OCT2* mutant brain (supplementary fig. 5), which may reflect a compensatory mechanism attenuating the impact of tryptophan deficiency on 5-HT content.

Notwithstanding the modest contribution of tryptophan supplementation to tryptophan and 5-HT concentrations in *OCT2* mutant brain, this supplementation improved several depression-related behaviors (i.e., sucrose preference, object location and coat state) in mutant mice, leading to similar performances to those of WT mice. This last observation underlines the importance of tryptophan availability for fluoxetine antidepressant efficacy, despite the mitigated effects of supplementation on 5-HT content in the mutants. Several mechanisms might be responsible for this beneficial effect of tryptophan supplementation on depressive-like behaviors. The firing of DRN 5-HT neurons is expected to facilitate antidepressant response [14], while this firing may also be affected by variations in the expression and/or the sensitivity of 5-HT1A autoreceptors located on these neurons [15,16]. The electrophysiological data presented here show that tryptophan supplementation did not alter significantly DRN 5-HT neuronal firing in OCT2 mutant mice, as could have been expected from its impact on behavior. However, this experiment led to an interesting observation, indicating that OCT2 activity by itself influences DRN firing during prolonged fluoxetine treatment. Paradoxically, SSRI resistance in *OCT2* mutant mice was accompanied with an increase (rather than a decrease) in DRN 5-HT neuron firing, suggesting that in our conditions firing activity was not the main mechanism influencing response to treatment. Such an uncoupling between DRN 5-HT neuron firing behavior and antidepressant response has previously been reported. 5-HT neurons in *Tph2*^-/-^ mice retain normal firing behavior despite the lack of brain 5-HT [68] and impaired response to chronic fluoxetine administration [26]. These earlier studies along with the present report support the idea that 5-HT deficiency may affect the response to SSRIs by curbing SSRI-induced 5-HT release independently of DRN neuron firing.

A well-known action of SSRI is the progressive desensitization of DRN 5-HT1A receptors, leading to increased firing of DRN 5-HT neuron and 5-HT release, a process that has been associated with the delay in therapeutic response and treatment resistance [15,69]. Other studies also suggest that tryptophan availability could modulate the tonic autoinhibition of neuronal firing in the DRN [70,71]. In the present study, DRN 5-HT neuron firing was relatively low after three-week fluoxetine treatment compared to corticosterone alone, indicating that the inhibition exerted by 5-HT1A autoreceptors is still potent. In these experimental conditions, these firing rates were significantly higher in *OCT2* mutant mice than in WT mice after fluoxetine treatment. Additionally, 5-HT1A autoreceptor *sensitivity* after corticosterone plus fluoxetine treatment did not significantly differ between genotypes (Fig. 3c), suggesting that these receptors should respond similarly to local 5-HT concentrations. Thus, slower desensitization of 5-HT1A autoreceptors and/or decreased 5-HT1A autoreceptors sensitivity can be excluded to explain the increase in DRN 5-HT neuronal activity in *OCT2* mutant mice, and dismissed as a main cause for their resistance to SSRI treatment. Hypothetically, decreased autoinhibition resulting from decreased extracellular 5-HT concentration in DRN of *OCT2* mutant mice compared to WT mice could account for the increased 5-HT neuron firing observed in these mutants, in fluoxetine-treated conditions.

Finally, to further refine the mechanisms underlying antidepressant resistance, we investigated the general consequences of the absence of OCT2 and tryptophan supplementation on intracellular signaling in the brain. The therapeutic effects of SSRI occurring after several weeks of treatment engage slow-onset, brain-wide, functional modifications of neural circuits, involving structural and synaptic plasticity. A number of these modifications converge on the limbic and prefrontal cortical areas of the brain, which regulate affect and cognitive processes, to drive the modulation of intracellular signaling cascades and of expression of target genes [72]. Specifically, alterations of MAP kinase ERK1/2, GSK3β, and mTOR signaling have been reported in the brain of depressed patients [49-51], and after antidepressant treatment in animal models [51-53]. Our findings show that the activation state of these three pathways in the dorsal hippocampus and prefrontal cortex responded vigorously to corticosterone treatment in parallel with the emergence of depression-related behavioral anomalies and were for most of them oppositely regulated by fluoxetine treatment, confirming previous observations [35]. These Western blot analyses pointed to anomalies in *OCT2* mutant mice brain of GSK3β and mTOR signaling, which might underlie resistance to fluoxetine treatment. Strikingly, tryptophan supplementation in *OCT2* mutant mice recruited selectively the mTOR protein complex 2. This activation may be involved in the partial rescue of the depression-like phenotype of *OCT2* mutant mice during fluoxetine treatment. mTOR signaling regulates key cellular functions and fundamental developmental and metabolic processes. The mTORC2 complex 2 specifically has been involved in hippocampal plasticity, learning and memory [73,74], and in the modulation of dopamine neurotransmission. *Rictor* null mice with disruption of neuronal mTORC2 show alterations in dopamine content and expression of dopamine receptors [75,76] and in basal or drug-induced locomotory activity [76]. Furthermore, mTORC2 invalidation specifically in dopaminergic areas was found to alter dopamine neuron structure and impair the rewarding response to morphine [77]. As the meso-limbic pathway plays an essential role in reward and motivation [78] we speculate that increased mTORC2 signaling in one or several regions of the brain could mediate the preferential effects of tryptophan supplementation on anhedonia in the *OCT2* mutants.

In conclusion, the present study provides the first evidence of the physiological relevance of OCT2 activity for tryptophan distribution in the brain, and its biological consequences on SSRI efficacy, mood and behavior. The interactions between dietary status and OCT2 transporter activity unveiled here may have important consequences in the field of depression therapy. In humans, variability in OCT2 activity may arise from genetic polymorphism [79], or by modulation of transporter activity by phosphorylation [80] or trafficking [81,82]. Moreover, a wide range of clinically used drugs, including antidiabetics and antiviral and antineoplastic agents, can interact with OCT [83]. It is conceivable that medications used to treat common pathologies could interfere with OCT2 transport across brain barrier structures. Whether and how these diverse genetic, epigenetic and drug interaction processes affecting OCT2 function may influence resistance to antidepressant treatment deserves to be carefully scrutinized. These findings also underscore the importance of unbalanced diets as potential risk factors for individual resistance to antidepressant treatment, and open the possibility to manipulate and correct antidepressant efficacy by dietary interventions in specific cases.

## ACKNOWLEDGEMENTS

We thank F. Machulka at the rodent phenotyping facility of Neuroscience Paris Seine for expert assistance with animal care and Jean-François Gilles at the imaging facility of Institut de Biologie Paris-Seine for help with immunofluorescence imaging. Financial support was provided by the Agence Nationale de la Recherche (ANR-13-SAMENTA-0003-01). The authors declare no conflict of interest.

## AUTHOR CONTRIBUTIONS

AO, BPG, JC, JML, VV and SG designed the experiments. AO, BPG, SM, JC, FL, AP, CG, CB and VV conducted the experiments and performed the data analysis. AO, BPG, JML, CB, VV and SG wrote the manuscript.

