## Supplementary information for "Organic cation transporter 2 contributes to SSRI antidepressant efficacy by controlling tryptophan availability in the brain"

### SUPPLEMENTARY MATERIAL AND METHODS

#### Behavioral tests

The coat state was assessed weekly as a measure of motivation toward self-care. It was evaluated as the sum of the score of different parts of the body, ranging between 0 for a well-groomed coat and 1 for an unkempt coat for head, neck, dorsal/ventral coat, tail, and forepaws/hind paws. For the sucrose preference test, singled-house mice were first habituated for 48 h to drink water from two bottles. On the following 3 days, the mice could choose between a water bottle and a 1% (wt/vol) sucrose solution bottle, with the position switched daily. Sucrose solution intake for 24 h was measured during the last 2 days and expressed as a percentage of the total amount of liquid ingested. The social interaction test was performed in a white open-field (42 x 42 cm) containing an empty wire mesh cage (10 x 6.5 cm) located at an extremity of the field in a low luminosity environment (25 lux). Individual mice were allowed to explore the open-field for two consecutive sessions of 2.5 min. During the second session, an unfamiliar mouse was introduced into the wire mesh cage. Between the two sessions, the test mouse was placed back into its home cage for approximately one minute. The time spent by the test mouse in the interaction zone, defined as an 8-cm-wide region surrounding the mesh cage, was measured in both sessions by video tracking (Viewpoint). For the object location test, the mice were habituated during two successive days to an open-field containing an intra-field cue (one wall covered with black and white stripes). Each mouse was allowed to freely explore the open-field for a 30-min period on day 1 and for two 10-min sessions separated by 5 h on day 2. On the third day, the test mouse was allowed to explore for 5 min two identical objects (5 x 2.5 cm) positioned in two adjacent corners of the open-field (acquisition phase) then returned to its home cage for 1 h. For the sample phase trial, one of the two objects was displaced to the opposite corner of the open-field. The time spent exploring both objects was recorded over a 5-min session by video tracking. The elevated O-maze consisted of an annular runway positioned 40 cm above the floor and divided into two opposing 90° closed sectors and two 90° open sectors. Mice were individually placed in the closed sector and their behavior recorded over a 5-min period. The time spent in each sector and the number of sector entries (a sector entry was defined as all four paws being placed in a sector) were determined by video tracking (Viewpoint, Lyon, France).

#### Western blotting

Samples were homogenized by sonication in 2 vol of ice-cold phosphate-buffered saline containing 1% Triton X-100, protease inhibitors (Complete Protease Inhibitor Cocktail, Roche Diagnostics, Meylan, France) and phosphatase inhibitors (Phosphatase Inhibitor Cocktail 3; Sigma-Aldrich, Darmstadt, Germany). Protein concentrations were determined by Bradford’s method. Protein samples (15 μg) suspended in NuPage LDS sample buffer (Invitrogen, Carlsbad, CA, USA) were separated by Bis-Tris sodium dodecyl sulfate polyacrylamide gel electrophoresis (10% gels) and transferred onto nitrocellulose membranes (Invitrogen). Transfer efficacy was controlled by Ponceau S staining. Unspecific binding sites were blocked with Tris-buffered saline containing 0.1% Tween-20 and 5% nonfat milk and membranes were immunoprobed with antibodies against Erk1/2 (1/1500, Cat# sc-135900, RRID:AB_2141283) from Santa Cruz Biotechnology (CA, USA), phosphorylated extracellular-signal regulated kinase1/2 (pErk1/2, 1/1000, Cat# 9101, RRID:AB_331646), p-P70S6 (1/500, Cat# 9206, RRID:AB_331768), Akt (1/2000, Cat# 2920, RRID:AB_1147620), pThr308 (1/200, # 4056, RRID:AB_331163) or Ser473 Akt (1/200, Cat# 9271, RRID:AB_329826) and phosphorylated GSK3β (p GSK3β, 1/1000, Cat# 9336, RRID:AB_331406) from Cell Signaling (Danvers, MA, USA), glycogen synthase kinase-3β (GSK3β, 1/2000, Cat# 4414, RRID:AB_259907) or β-actin (1/2500; Cat# 4700, RRID:AB_476730) from Sigma-Aldrich. Membranes were incubated with infrared-labeled secondary antibodies (IRDye 700DX RRID:AB_220144 and IRDye 800CW, RRID:AB_220150; 1/5000) from Rockland (Gilbertsvillle, PA, USA). Immunoblotting was quantified with an Odyssey Infrared Imaging System and Application Software version 3.0 (LI-COR Biosciences, Lincoln, NE, USA).

### SUPPLEMENTARY RESULTS

**Supplementary table 1 Statistical analysis of behavior and coat state (Figure 1)**

Two-way ANOVA (n = 10-14) followed by Tukey's post hoc test for behavior and Dunnett's for coat state.

Main effects of tryptophan supplementation, genotype and/or genotype*tryptophan supplementation interaction

| Test | Significance |
|---|---|
| Sucrose preference | Treatment $F_{2, 74}$ = 22.43; P < 0.0001<br>Genotype $F_{1, 74}$ = 4.999; P = 0.0284<br>Interaction $F_{2, 74}$ = 8.820; P = 0.0004 |
| Social interaction | Treatment $F_{2, 60}$ = 9.377; P < 0.0003<br>Genotype $F_{1, 60}$ = 1.204; P = 0.2769<br>Interaction $F_{2, 60}$ = 2.642; P = 0.0795 |
| Object location | Treatment $F_{2, 56}$ = 3.863; P = 0.0268<br>Genotype $F_{1, 56}$ = 11.09; P = 0.0015<br>Interaction $F_{2, 56}$= 2.886; P = 0.0642 |
| Elevated O-maze | Treatment $F_{2, 61}$ = 43.30; P < 0.0001<br>Genotype $F_{1, 61}$ = 0.3589; P=0.5513<br>Interaction $F_{2, 61}$ = 5.737; P = 0.0052 |
| Coat state | Treatment $F_{10, 278}$= 59.78; P < 0.0001<br>Genotype $F_{1, 278}$ = 8.384; P < 0.0041<br>Interaction $F_{10, 278}$= 1.101; P=0.3610 |

**Supplementary table 2 Statistical analysis of behavior and coat state after tryptophan supplementation (Figure 3A-D)**

Two-way ANOVA (n = 5-8) followed by Tukey's post hoc test for behavior and Dunnett's for coat state.

Main effects of treatment, genotype and/or genotype*treatment interaction

| Test | Significance |
|---|---|
| Sucrose preference | Treatment $F_{3, 72}$= 16.54; P < 0.0001<br>Genotype $F_{1, 72}$= 2.297; P = 0.1340<br>Interaction $F_{3, 72}$= 3.142; P = 0.0304 |
| Social interaction | Treatment $F_{3, 57}$= 10.63; P < 0.0001<br>Genotype $F_{1, 57}$ = 0.4529; P = 0.5037<br>Interaction $F_{3, 57}$= 3.142; P = 0.0321 |
| Object location | Treatment $F_{2, 38}$ = 15.14; P<0.0001<br>Genotype $F_{1, 38}$ = 1.901; P=0.1760<br>Interaction $F_{2, 38}$= 2.568; P = 0.0899 |
| Coat state | Fluoxetine treatment 8,176 = 56.27; P < 0.0001<br>Genotype $F_{3, 22}$= 1.122; P = 0.3616<br>Interaction $F_{24, 176}$= 3.151; P < 0.0001 |

**Supplementary table 3 Statistical analysis of HPLC data after tryptophan supplementation (Figure 4)**

Two-way ANOVA (n = 5-7) followed by Sidak's post hoc analysis of the effect of tryptophan supplementation.

Main effects of tryptophan supplementation, genotype and/or genotype*tryptophan supplementation interaction

| **Condition Figure 4A** | **Significance** |
|---|---|
| Tryptophan/Plasma | Supplementation F1,19 = 52.22; P < 0.0001<br>Genotype F1, 19 = 46.89; P < 0.0001<br>Interaction F1, 19 = 0.03468; P = 0.8542 |
| Kynurenin/Plasma | Supplementation F1,19 = 38.29; P < 0.0001<br>Genotype F1,19 = 44.64; P < 0.0001<br>Interaction F1, 19 = 3.124; P = 0.0932 |

| **Condition Figure 4B** | **Significance** |
|---|---|
| Tryptophan/striatum | Supplementation F1,20 = 26.63; P < 0.0001<br>Genotype F1, 20 = 116.9; P < 0.0001<br>Interaction F1, 20 = 2.271; P = 0.1474 |
| Tryptophan/hippocampus | Supplementation F1,20 = 9.245; P = 0.0065<br>Genotype F1,20 = 78.78; P < 0.0001<br>Interaction F1,20 = 4.455; P = 0.0476 |
| Tryptophan/cortex | Supplementation F1,19 = 21.50; P = 0.0002<br>Genotype F1,19 = 110.1; P < 0.0001<br>Interaction F1,19 = 1.909; P = 0.1831 |
| 5-HT/striatum | Supplementation F1,20 = 11.61; P = 0.0028<br>Genotype F1,20 = 8.757; P = 0.0078<br>Interaction F1,20 = 0.002546; P = 0.9603 |
| 5-HT/hippocampus | Supplementation F1,20 = 9.335; P = 0.0062<br>Genotype F1,20 = 3.068; P = 0.0952<br>Interaction F1,20 = 1.811; P = 0.1935 |
| 5-HT/cortex | Supplementation F1,19 = 1.608; P = 0.2200<br>Genotype F1,19 = 7.250; P=0.0144<br>Interaction F1,19 = 0.2478; P = 0.6243 |
| Kynurenine/striatum | Supplementation F1,20 = 5.066; P = 0.0358<br>Genotype F1,20 = 79.33; P < 0.0001<br>Interaction F1,20 = 16.63; P = 0.0006 |
| Kynurenine/hippocampus | Supplementation F1,20 = 1.440; P = 0.2442<br>Genotype F1,20 = 13.35; P = 0.0016<br>Interaction F1,20 = 3.134; P = 0.0919 |
| Kynurenine/cortex | Supplementation F1,19 = 7.126; P = 0.0152<br>Genotype F1,19 = 30.96; P < 0.0001<br>Interaction F1,19 = 0.2656; P = 0.6122 |

**Supplementary table 4 Detailed data of electrophysiological recordings of DR 5-HT neurons**

| | CORT | CORT/FLX | CORT/FLX/TRP |
|---|---|---|---|
| OCT2 +/+ | 3.02 ± 0.21 (n= 65; 7 mice) | 0.72 ± 0.09 (n= 54; 7 mice) | 1.02 ± 0.13 (n= 34; 5 mice) |
| OCT2 -/- | 2.34 ± 0.17 (n= 45; 5 mice) | 1.28 ± 0.19 (n= 45; 4 mice) | 1.28 ± 0.14 (n= 60; 5 mice) |

**Supplementary table 5 Statistical analysis of Western blot data (Figure 5A)**

Two-way ANOVA (n = 6-9) followed by Tukey's post hoc test.

Main effects of treatment, genotype and/or genotype*treatment interaction

| Conditions | Significance |
|---|---|
| pERK1/2/ERK1/2 (% control)<br>Hippocampus | Treatment $F_{2, 32}$= 87.20; $P < 0.0001$<br>Genotype $F_{1, 32}$ = 5.995; $P = 0.0200$<br>Interaction $F_{2, 32}$= 0.8528; $P = 0.4357$ |
| pERK1/2/ERK1/2 (% control)<br>Cortex | Treatment $F_{2, 34}$= 78.72; $P < 0.0001$<br>Genotype $F_{1, 34}$= 0.008852; $P = 0.9256$<br>Interaction $F_{2, 34}$= 0.02774; $P = 0.9727$ |
| pThr389-P70S6K/β-actin (% control)<br>Hippocampus | Treatment $F_{2, 37}$= 16.05; $P < 0.0001$<br>Genotype $F_{1, 37}$= 0.1084; $P = 0.7439$<br>Interaction $F_{2, 37}$= 1.787; $P = 0.1815$ |
| pThr389-P70S6K/β-actin (% control)<br>Cortex | Treatment $F_{2, 37}$= 20.12; $P < 0.0001$<br>Genotype $F_{1,37}$ = 0.001220; $P = 0.9723$<br>Interaction $F_{2, 37}$= 0.01419; $P = 0.9859$ |
| pSer473-Akt/Akt (% control)<br>Hippocampus | Treatment $F_{2, 37}$= 12.06; $P < 0.0001$<br>Genotype $F_{1, 37}$= 2.662; $P = 0.1113$<br>Interaction $F_{2, 37}$= 4.595; $P = 0.0165$ |
| pSer473-Akt/Akt (% control)<br>Cortex | Treatment $F_{2, 39}$= 21.75; $P < 0.0001$<br>Genotype $F_{1,39}$ = 0.07799; $P = 0.7815$<br>Interaction $F_{2, 39}$= 1.896; $P = 0.1637$ |
| pSer9-GSK3β/GSK3β (% control)<br>Hippocampus | Treatment $F_{2, 31}$= 21.14; $P < 0.0001$<br>Genotype $F_{1,31}$ = 1.816; $P = 0.1876$<br>Interaction $F_{2, 31}$= 17.11; $P < 0.0001$ |
| pSer9-GSK3β/GSK3β (% control)<br>Cortex | Treatment $F_{2, 35}$= 22.43; $P < 0.0001$<br>Genotype $F_{1,35}$ = 0.006246; $P = 0.9375$<br>Interaction $F_{2, 35}$= 0.1075; $P$=0.8984 |

### SUPPLEMENTARY FIGURES

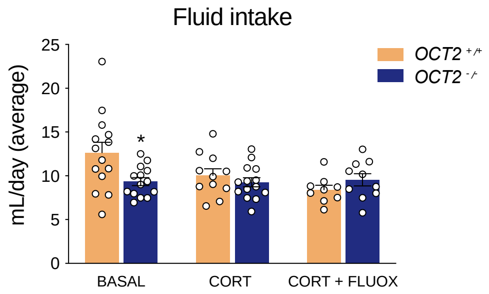


**Supplementary figure 1. Total fluid intake during the sucrose preference test (average over 3 days) at basal state (BASAL), after corticosterone treatment (CORT), and after 3 weeks of fluoxetine plus corticosterone (CORT/FLUOX).** Two-way ANOVA analysis (n =10–14) shows a main effect of treatment ($F_{2,66}$ = 3.529; $P = 0.0350$) but not genotype ($F_{1,66}$ = 2.464; $P = 0.1213$). WT (*OCT2*$^{+/+}$) and *OCT2* mutant (*OCT2*$^{-/-}$) mice drink similar amounts of fluid after corticosterone and corticosterone plus fluoxetine treatment. Results are given as mean ± s.e.m.

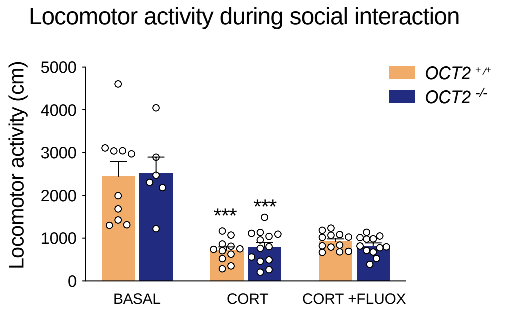


**Supplementary figure 2. Locomotor activity during social interaction at basal state (BASAL), after corticosterone treatment (CORT), and after 3 weeks of fluoxetine plus corticosterone (CORT/FLUOX) in WT (*$OCT2^{+/+}$*) and *OCT2* mutant (*$OCT2^{-/-}$* ).** Two-way ANOVA analysis (n = 10–14) shows a main effect of treatment (F2,58 = 50.61; P < 0.0001) but not genotype (F1,58 = 0.0116; P = 0.9146). Results are given as mean ± s.e.m.

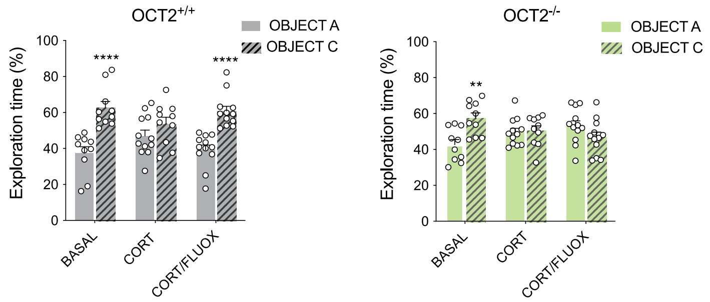


**Supplementary figure 3. Object location test at basal state (BASAL), after corticosterone treatment (CORT), and after 3 weeks of fluoxetine plus corticosterone (CORT/FLUOX) in WT (*$OCT2^{+/+}$*) and *OCT2* mutant (*$OCT2^{-/-}$* ).** Two-way ANOVA analysis (n = 10–14) shows a main effect of object (F1,60 = 45.55; P < 0.0001) and interaction object*genotype (F2,60 = 4.547; P = 0.0145) but not genotype (F2,60 = 0.005847; P = 0.9942) for WT mice (*$OCT2^{+/+}$*); and interaction object*genotype (F2,60 = 7.553; P = 0.0012) but not treatment (F2,60 = 0.0001404; P = 0.999) or object (F1,60 = 1.936; P = 0.1692) for *OCT2* mutant mice (*$OCT2^{-/-}$*). Sidak's post hoc, object A versus object C, *** P < 0.0001.

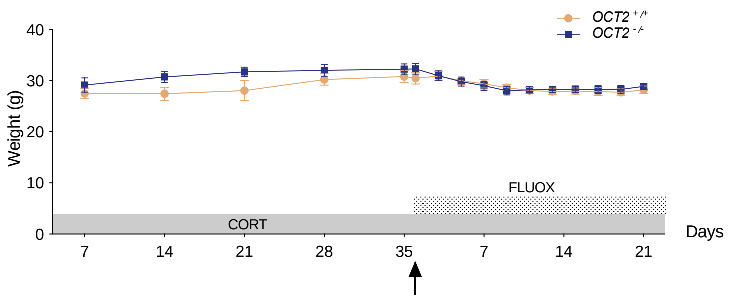


**Supplementary figure 4. Weight of WT (*$OCT2^{+/+}$*) or *OCT2* mutant mice (*$OCT2^{-/-}$*) during corticosterone and fluoxetine treatment.** Two-way ANOVA analysis (n = 10–14) show a main effect of time of treatment (F15,405 = 3.566; <0.0001) and of genotype (F1,405 = 7.046; P = 0.0083) but not time*genotype interaction (F15, 405 = 0.8053; P = 0.6720).

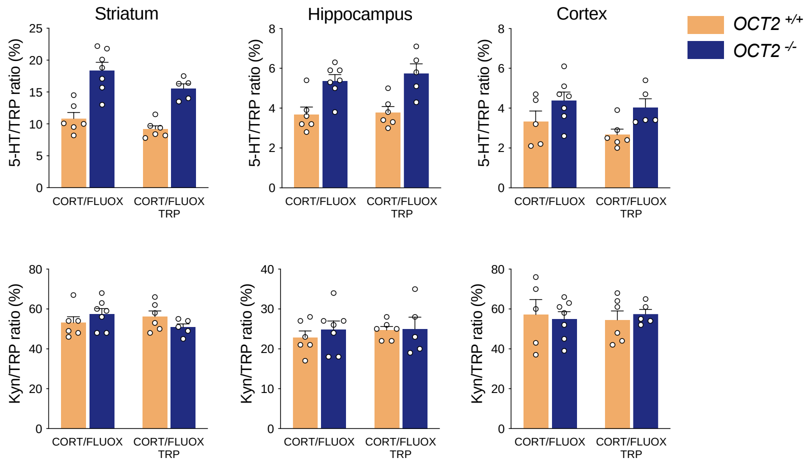


**Supplementary figure 5. 5-HT/tryptophan and kynurenine/tryptophan ratios in brain of WT (*OCT2*$^{+/+}$) or *OCT2* mutant mice (*OCT2*$^{-/-}$) during corticosterone and fluoxetine treatment with or without tryptophan supplementation.** Two-way ANOVA analysis (n = 10–14) show a main effect of genotype in striatum (F1,20 = 49.62; P <0.0001), hippocampus (F1,20 = 25.02; P = 0.0001) and (F1,19 = 8.262; P = 0.0097), reflecting increased TPH2 activity. Kynurenine/tryptophan ratios ratios were comparable in all groups, suggesting no effect of genotype or of tryptophan supplementation on kynurenine production.

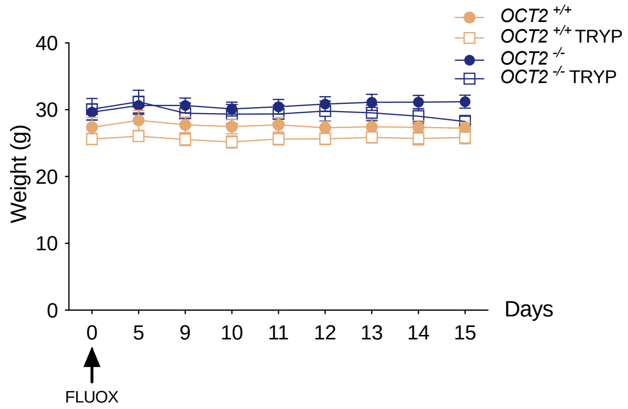


**Supplementary figure 6. Weight of WT (*OCT2*$^{+/+}$) or *OCT2* mutant mice (*OCT2*$^{-/-}$) during corticosterone plus fluoxetine treatment with (TRP) or without tryptophan supplementation.** Two-way ANOVA analysis (n = 5–6) shows no main effect of time in WT (F1.956,23.47 = 1.655; P = 0.2128) or group (F1.872,16.85 = 1.810; P = 0.1951) mice. Results are given as mean ± s.e.m.

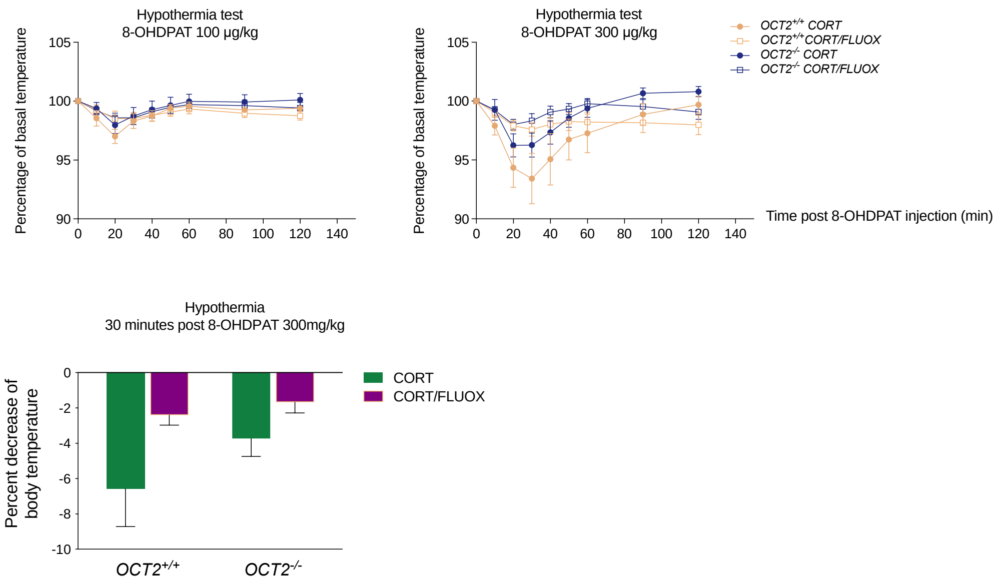


**Supplementary figure 7. Kinetics of 8-OH-DPAT-induced hypothermia and temperature change in WT (*OCT2*$^{+/+}$) or *OCT2* mutant mice (*OCT2*$^{-/-}$).** Two-way ANOVA analysis (n = 8–10) shows a main effect of group ($F_{2.138,70.56} = 20.86$; $P < 0.0001$), and of interaction group* time ($F_{24,64} = 3.444$; $P < 0.0001$), but no main effect of time ($F_{3,33} = 1.348$; $P = 0.2758$) on percentage of basal temperature at the dose of 300 µg/kg; and a main effect of treatment ($F_{1,33} = 6.206$; $P = 0.0179$) but not of genotype ($F_{1,33} = 2.046$; $P = 0.1620$) or interaction ($F_{1,33} = 0.718$; $P = 0.403$) on percentage of decrease in temperature. Results are given as mean ± s.e.m.
